# On the analysis of functional PET (fPET)-FDG: baseline mischaracterization can introduce artifactual metabolic (de)activations

**DOI:** 10.1101/2024.10.17.618550

**Authors:** Sean E. Coursey, Joseph Mandeville, Murray B. Reed, Grant A. Hartung, Arun Garimella, Hasan Sari, Rupert Lanzenberger, Julie C. Price, Jonathan R. Polimeni, Douglas N. Greve, Andreas Hahn, Jingyuan E. Chen

## Abstract

Functional Positron Emission Tomography (fPET) with (bolus plus) constant infusion of [^18^F]-fluorodeoxyglucose FDG), known as fPET-FDG, is a recently introduced technique in human neuroimaging, enabling the detection of dynamic glucose metabolism changes within a single scan. However, the statistical analysis of fPET-FDG data remains challenging because its signal and noise characteristics differ from both classic bolus-administration FDG PET and from functional Magnetic Resonance Imaging (fMRI), which together compose the primary sources of inspiration for analytical methods used by fPET-FDG researchers. In this study, we present an investigate of how inaccuracies in modeling baseline FDG uptake can introduce artifactual patterns to detrended TAC residuals, potentially introducing spurious (de)activations to general linear model (GLM) analyses. By combining simulations and empirical data from both constant infusion and bolus-plus-constant infusion protocols, we evaluate the effects of various baseline modeling methods, including polynomial detrending, regression against the global mean time-activity curve, and two analytical methods based on tissue compartment model kinetics. Our findings indicate that improper baseline removal can introduce statistically significant artifactual effects, although these effects characterized in this study (∼2-8%) are generally smaller than those reported by previous literature employing robust sensory stimulation (∼10-30%). We discuss potential strategies to mitigate this issue, including informed baseline modeling, optimized tracer administration protocols, and careful experimental design. These insights aim to enhance the reliability of fPET-FDG in capturing true metabolic dynamics in neuroimaging research.

## 1 Introduction

Functional Positron Emission Tomography (fPET) using (bolus plus) continuous infusion of [^18^F]-fluorodeoxyglucose (FDG) is a recent advancement in human neuroimaging that allows for the tracking of minute-scale dynamic changes in glucose metabolism within a single scan (Villien et al., 2014). Previously, continuous infusion of PET radiotracers has been practiced since the 1990s to enable more accurate receptor-binding measures and the tracking of dynamic changes in neurotransmitter concentrations (Carson et al., 1993; Sander et al., 2020) and continuous-infusion FDG scanning has been shown, in small animal studies, to be effective at identifying transient changes in glucose metabolism (Bérard et al., 2006; Cauchon et al., 2012). However, it was not until 2014 that this approach was applied to functional metabolic imaging in humans (Villien et al., 2014). Villien et al. infused the FDG tracer at a constant rate, allowing its plasma concentration of FDG to slowly equilibrate to a steady level. This introduced a quasi-linear regime in the time-activity curve (TAC), where the slope is proportional to the cerebral metabolic rate of glucose consumption (CMRglu). Their simulations indicated that transient changes in glucose uptake introduce nonlinear deviations from this slope, while sustained changes can alter the equilibrium slope.

Since its inception in 2014, human fPET-FDG imaging has rapidly garnered attention, as it enables the measurement of dynamic changes in brain metabolism subserving brain computing across broad task and consciousness conditions (Hahn et al., 2020; Jamadar et al., 2019; Rischka et al., 2018; Stiernman et al., 2021). Emerging evidence suggests that fPET-FDG can improve temporal resolution by an order of magnitude compared to traditional bolus-injection FDG-PET (Hahn et al., 2024; Rischka et al., 2018). Moreover, its intra-scan dynamic nature also complements the hemodynamic information tracked by functional Magnetic Resonance Imaging (fMRI) (Godbersen et al., 2023; Jamadar et al., 2021; Stiernman et al., 2021; Verger and Guedj, 2018; Zürcher et al., 2024).

While the efficacy of fPET-FDG has been demonstrated by several studies that successfully captured metabolic (de)activations across various sensory and cognitive paradigms, the statistical analysis of fPET-FDG data remains an evolving field. Because, like fMRI, fPET-FDG also tracks intra-scan contrasts, existing fPET-FDG studies have often adopted statistical inference methods from the field of fMRI analysis, including the general linear model (GLM) framework for mapping task (de)activations (Hahn et al., 2016; Jamadar et al., 2019; Stiernman et al., 2021; Villien et al., 2014). However, differences in signal and noise characteristics between fPET-FDG and fMRI present new challenges when adapting these established statistical frameworks.

In a continuous-infusion paradigm, FDG accumulates in cells as they attempt to metabolize it over the course of a scan resulting in a monotonically increasing trend at baseline metabolism after decay correction (Villien et al., 2014). Unlike baseline drifts occurring in fMRI data, which exhibit much slower frequencies than the seconds-scale hemodynamic fluctuations of interest (Caballero-Gaudes and Reynolds, 2017; Chen and Glover, 2015), the fPET-FDG baseline comprises frequencies similar to the minutes-scale metabolic dynamics of interest, making it challenging to separate from task-induced effects.

Moreover, identifying appropriate reference regions for baseline correction is problematic because glucose metabolism occurs throughout the whole brain due to widespread neuronal and glial activity. Existing studies have employed various methods to isolate stimulus-driven metabolic changes from the baseline trend, including low-order polynomial detrending fit voxel-wise (Villien et al., 2014) or fit to the mean TAC and then scaled voxel-wise (Hahn et al., 2016; Khattar et al., 2024; Rischka et al., 2018; Stiernman et al., 2021), regressing against a task-independent TAC obtained by averaging task-free gray matter (Rischka et al., 2018), or simple global signal regression in the context of independent component analysis (Li et al., 2020).

However, if the shape of the baseline TACs is inaccurately characterized, this may introduce artifactual patterns into the detrended data’s time courses. In a GLM analysis, the task regressor can fit to the un-modeled portion of the baseline, biasing fitted coefficients and introducing artifactual shifts in the t-scores of the task effect. These artifacts may manifest as global shifts, from consistent mischaracterization, and/or anatomically coherent spatial patterns, due to structured information embedded in regionally varying tracer kinetics (Heiss et al., 1984; Volpi et al., 2023).

In this study, we combine simulations and empirical fPET-FDG data to investigate how various baseline modeling methods affect the TAC and whether mischaracterization can result in artifactual (de)activations. Specifically, we evaluate polynomial detrending, regression against the global mean TAC, and two analytical methods based on the tissue compartment model (Phelps et al., 1979). We first characterize the shapes of residual TACs left by various detrending methods and evaluate the extent to which these detrending approaches mischaracterize the actual baseline signal in both constant-infusion (CI) and bolus-plus-constant-infusion (B+CI) fPET-FDG. We then demonstrate that baseline mischaracterization can result in spatially structured artifactual (de)activations when the mischaracterized portion of baseline TAC exhibits certain degrees of correlation with the task, using an illustrative constant-infusion resting-state dataset expected to follow the null hypothesis of no task activation. Finally, we discuss potential strategies to mitigate the effects of baseline mischaracterization, including informed baseline modeling, optimized tracer administration protocols, and careful experimental design. These results emphasize the importance of accurate baseline modeling to statistical inference in fPET-FDG studies.

## 2 Methods

### 2.1 Empirical data

Two imaging datasets were employed to evaluate the influence of baseline mischaracterization on the statistical inference of fPET-FDG:

1. *Constant infusion (CI) dataset*: A publicly available resting-state fPET-fMRI dataset collected from healthy young adults (*N* = 27, 18-23 yrs, 21 female) by research groups at Monash University (Jamadar et al., 2020). The experiments were performed on a Siemens (Siemens Healthineers, Erlangen, Germany) 3 Tesla Biograph mMR scanner. Subjects were instructed to rest with their eyes open during the scan, which lasted approximately 95 minutes. [^18^F]FDG (average dose ∼6.3 mCi) was infused at a constant rate throughout the scan. The PET data were reconstructed with a nominal voxel size of 2.09 × 2.09 × 2.03 mm³, and the temporal frame length was 16 seconds. Of the 27 subjects, we removed three for data quality concerns (two for motion artifacts and one for an infusion pump error), resulting in *N* = 24 for our analyses.
2. *Bolus-plus-constant infusion (B+CI) dataset*: Simultaneous fPET-fMRI datasets collected at the Athinoula A. Martinos Center for Biomedical Imaging at Massachusetts General Hospital (MGH) from healthy young adults participating in two different experiments: a working-memory task (*N* = 11, total scan duration = 105 ± 0.3 minutes, mean and standard deviation across subjects) and an endogenous arousal study (*N* = 18, total scan duration = 91 ± 8 minutes, mean and standard deviation across subjects) (Figure S1A). All experimental procedures performed at MGH were approved by the Institutional Review Board, and all participants provided written informed consent. In the working-memory experiment, subjects performed letter-based two-back tasks (Chen et al., 2015) with jittered stimulus timings (Figure S1B). In the endogenous arousal experiment, subjects rested with eyes closed, and their within-scan arousal states were inferred from concurrent EEG or behavioral measures (Figure S1C). All experiments were conducted on a Siemens 3 Tesla Tim MAGNETOM Trio MR scanner (Siemens) with a BrainPET insert (Siemens). [^18^F]FDG was administered using a B+CI protocol (total dose ∼10 mCi, and the bolus dose was 20% of the constant infusion dose) (Rischka et al., 2018).

The PET data were first sorted into line-of-response space and subsequently compressed into sinogram space for reconstruction. A standard 3D ordinary Poisson ordered-subset expectation maximization algorithm was applied, incorporating both prompt and variance-reduced random coincidence events. The process also included normalization, scatter, and attenuation sinograms (Villien et al., 2014). Attenuation correction was performed using a pseudo-CT map derived from high-resolution anatomical MRI data (Izquierdo-Garcia et al., 2014). The PET data were reconstructed with a nominal voxel size of 2.5 × 2.5 × 2.5 mm³, and the temporal frame length was 30 seconds.

The jittered working-memory and the endogenous-arousal datasets were combined to characterize the artifactually patterned residuals left by various detrending methods on bolus-plus-constant infusion datasets. We anticipate that the jittering of the working-memory dataset and the natural variation in arousal timing of the endogenous arousal dataset cancel out systemic task and arousal-based effects at the group level (see Section 2.4).

### 2.2 Simulations

Simulations were performed to assess the residual fPET TAC patterns after detrending, which complement our empirical data analyses by allowing detrending methods to be tested on noise-free time-series data with known, well-defined signal characteristics. We used the irreversible two-compartment model to generate task-free, resting-state FDG TACs (Phelps et al., 1979). We actualized the model as a simulation using an Implicit Euler implementation. Briefly, the irreversible two-compartment model describes the transport of FDG from arterial blood into tissue (first compartment) and its phosphorylation within cells, where it becomes trapped as FDG-6-phosphate (second compartment). The model assumes reversible exchange between the blood and tissue compartments but irreversible phosphorylation. Using the model to simulate FDG TACs requires an input function of the arterial tracer concentration at each timepoint and kinetic constants describing the rate that the tracer is transported from the blood to the tissue (K_1_), back from the tissue to the blood (k_2_), and that the tracer is phosphorylated in tissue (k_3_). We referred to previous literature (Villien et al., 2014) for these rate constants (K_1_ = 0.1 mL/min/g, k_2_ = 0.15 min⁻¹, k_3_ = 0.08 min⁻¹).

Arterial input functions (AIFs) were extracted from the empirical data using an image-derived approach described previously (Sari et al., 2017) plus a model-fitting step to reduce noise. For the CI case, AIFs were extracted from the resting-state dataset; for the B+CI case, from the working-memory dataset. Carotid arteries were manually segmented for each subject from their high-resolution T1-weighted anatomical images and image-derived input functions (IDIFs) were extracted from the scans, and then smoothed AIFs were generated by fitting the convolution of a decaying biexponential response function with the infusion paradigm (Villien et al., 2014) to the IDIFs. The AIFs were then averaged to create group-level AIFs.

### 2.3 Preprocessing and cerebral cortical parcellation

For the CI dataset, motion correction was performed as described in a previous publication (Jamadar et al., 2020). For the B+CI data collected at MGH, all fPET images were motion-corrected by coregistering to the middle temporal frame using the 3dvolreg function in AFNI (Cox, 1996).

For voxel-wise analysis, individual fPET data were smoothed within a cerebral cortical gray-matter mask using an 8-mm full-width at half-maximum (FWHM) Gaussian kernel. For parcel-wise analysis, cortical gray-matter regions were delineated using the “7-network” parcellation from an existing multi-resolution functional atlas (Schaefer et al., 2018), examining the 100-, 300-, and 500-parcel resolutions. After delineating all brain parcels using the high-resolution anatomical data, the parcels were registered to the native image space of each individual’s fPET-FDG data to characterize region-specific metabolic patterns, using FreeSurfer (Fischl, 2012). No spatial smoothing was performed prior to extracting the mean signal of each parcel.

### 2.4 Characterizing residuals after baseline modeling

To characterize the shapes of residual TACs after detrending, we evaluated five baseline modeling methods:

- Third-order polynomial (P3): Regressing a third-order polynomial fitted to the time series of each voxel or parcel.
- Third-order polynomial pre-fitted to mean TAC (P3MT): Fitting a third-order polynomial to the mean gray-matter TAC and then regressing it from each voxel or parcel.
- Mean TAC (MT): Regressing the mean cerebral cortex gray-matter TAC from each voxel or parcel time series.
- Spectral analysis (SA): Modeling the baseline using components identified via spectral analysis, requiring prior knowledge of the AIF (Cunningham and Jones, 1993).
- Linear plus bi-exponential model (EXP2): Fitting a linear plus biexponential function based on the tissue compartmental model (see Appendix).

Note that in addition to the low-order polynomial and mean TAC methods used in existing fPET-FDG studies (Hahn et al., 2016; Li et al., 2020; Stiernman et al., 2021; Villien et al., 2014), we also tested two novel analytical solutions of baseline models (“SA” and “EXP2”). The AIFs for spectral analysis were created as described in Section 2.2 from the resting-state dataset for the constant-infusion analyses and the working memory dataset for the bolus-plus-constant-infusion analyses. All methods were applied both including and excluding the initial 10 minutes of the TAC to examine the impact of the initial dynamic phase before tracer equilibration (Villien et al., 2014).

These detrending methods were applied to both the empirical resting-state CI data and the composite B+CI datasets. To rule out true effects as the primary driver of residual patterning, we applied the same detrending methods to simulated resting-state CI and B+CI TACs (see Section 2.2) and compared the results to those derived from the empirical data. Assuming the dataset follow the null hypothesis of no true effect, which we expect them to, perfect detrending would result in a flat residual time-series with the only deviations from zero being noise, any consistent deviations from zero must therefore be attributed to imperfect detrending.

### 2.5 Assessing spurious metabolic (de)activations in GLM-based statistical inference

To assess whether baseline mischaracterization could introduce structured artifactual (de)activations, we applied a series of sham task regressors to the empirical fPET-FDG data and evaluated the GLM-based statistical results at the group level, using the resting-state CI dataset as an illustrative example. Although we also analyzed the group-level artifactual effects in the composite B+CI data (see Figures S2 and S3), we chose the CI data for demonstration because its inter-subject variability reflects that of a typical study (i.e., all subjects underwent identical experiment procedures). In contrast, the B+CI dataset, being a composite of working memory and endogenous arousal data, may exhibit higher-than-normal inter-subject variability due to the additional between-experiment differences, complicating interpretation. Additionally, while the task and arousal effects may be negligible in the group-mean TAC due to jitter and randomness, allowing us to evaluate the residual patterns from different detrending methods (Section 2.4), the true task and arousal effects remain present at the single-subject level.

The sham task paradigms consisted of alternating 10-minute “on” and “off” blocks with varying initial rest periods, resulting in regressors with different extents of correlation with the residual time-series patterns left after baseline removal. Three regressors were selected to represent high positive correlation, negligible correlation, and moderate negative correlation with the mean residual pattern of P3MT, using this detrending method as an exemplar. The 10-minute block durations are comparable to those used in previous fPET studies (Hahn et al., 2016; Kraus et al., 2020; Li et al., 2020; Villien et al., 2014). Each sham regressor was applied in a GLM analysis along with the baseline TAC model, using each of the five baseline modeling methods, both including and excluding the initial 10 minutes of data. For the P3MT and SA analyses, the task regressor was included as a nuisance when creating baseline regressors from the mean gray-matter TAC.

To evaluate the impact of spatial smoothing on artifactual effects, GLM analyses were performed at various levels of applied smoothing: voxel-wise with 8-mm FWHM smoothing, and at the 500-, 300-, and 100-parcel levels.

To evaluate spurious metabolic (de)activations introduced by each analysis for both significance and effect size, we computed both the random-effects t-scores and the average percent signal change (PSC) across subjects. The PSC, a proxy for the percent change in CMRglu associated with a task, is calculated by comparing the slope of the “on” portion of the fitted task regressor to the slope of the latter portion of the fitted baseline (Godbersen et al., 2024).

## 3 Results

### 3.1 Residual patterns after baseline modeling

Figure 1 summarizes the residual patterns after applying different baseline removal methods to both empirical and simulated data. Polynomial detrending (“P3” and “P3MT”) introduced a distinct pattern in the mean residual TACs for both resting-state CI and composite B+CI data. Regression against the global mean TAC, by construction, did not produce a mean residual pattern at the whole-brain level, but it did introduce regional residual patterns evidenced by the different patterns emerging from different ROIs (e.g. “MT ROI 1” and “MT ROI 2”, showing the most pronounced cases). While P3MT produced a nearly identical mean residual time-series pattern to P3 at the whole-brain level, the spatial distribution of the magnitude of the pattern by region bore some resemblance to that of MT, which can be seen in Figure 2.

**Figure 1:**
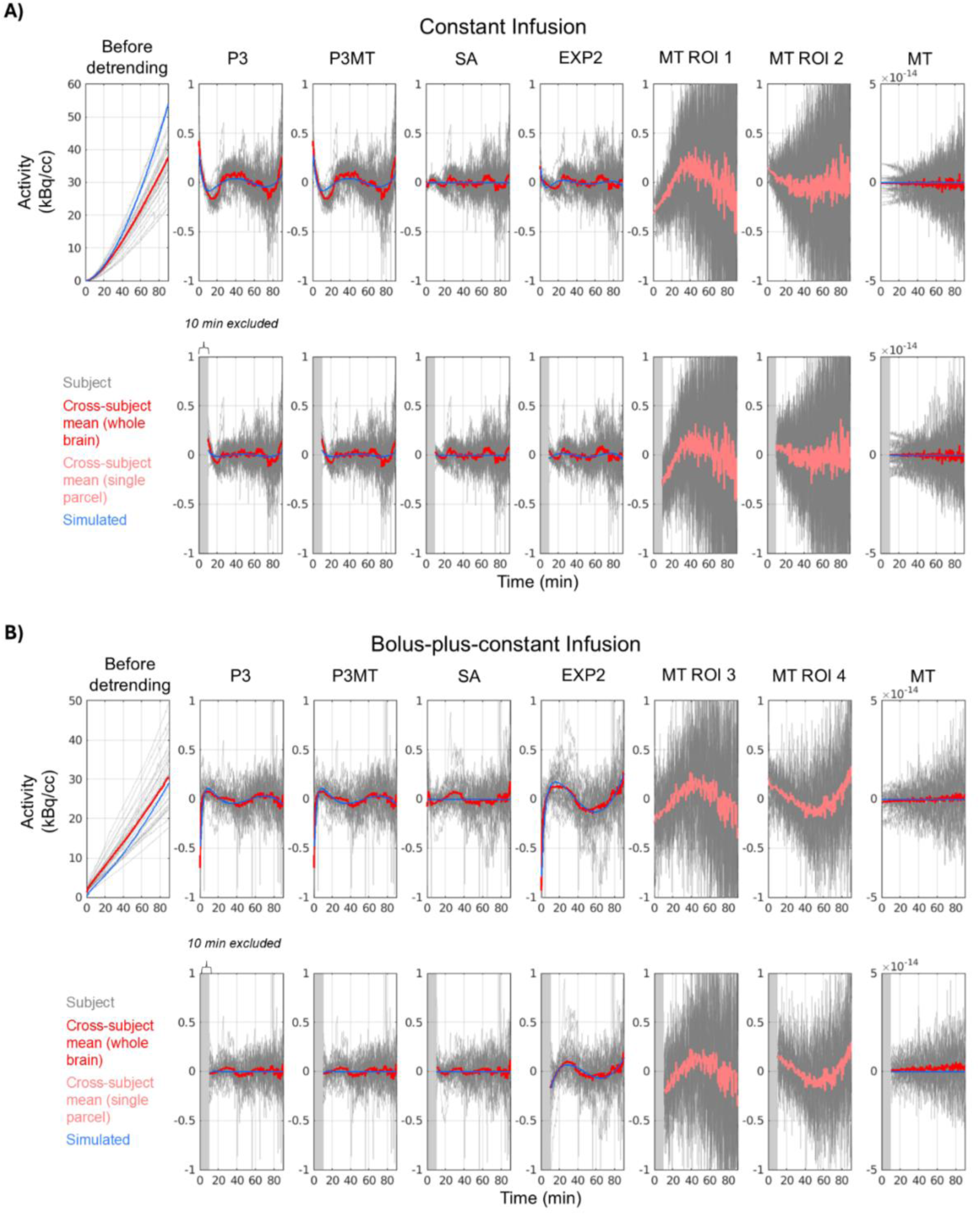
Residuals from CI (**A**) and B+CI (**B**) data and simulations after applying various detrending methods. Whole-brain TACs are plotted with a red line indicating the cross-subject mean; TACs from a single parcel are plotted with a pink line indicating the cross-subject mean. Gray blocks indicate excluded time-points. Blue indicates simulated resting-state TACs. Third-order polynomial detrending, whether applied parcel-wise (P3) or pre-fitted to the mean TAC (P3MT), leaves a consistent residual pattern across the brain for both CI and B+CI data. Excluding the first 10 minutes decreases this pattern, especially for data with a bolus. Regression of the global mean TAC (MT) leaves residual patterns in individual parcels (e.g. ROIs 1–4, being parcels 29, 77, 97, and 52 in Fig. 2, respectively, chosen for their large residual patterns, i.e., representing a potential worst-case scenario), but, by construction, leaves no average pattern across the brain. Spectral analysis (SA) leaves a smaller residual pattern than polynomial detrending for both infusion paradigms, while a linear plus bi-exponential model (EXP2) performs comparably to SA for the CI data but performs poorly for the B+CI data.

**Figure 2:**
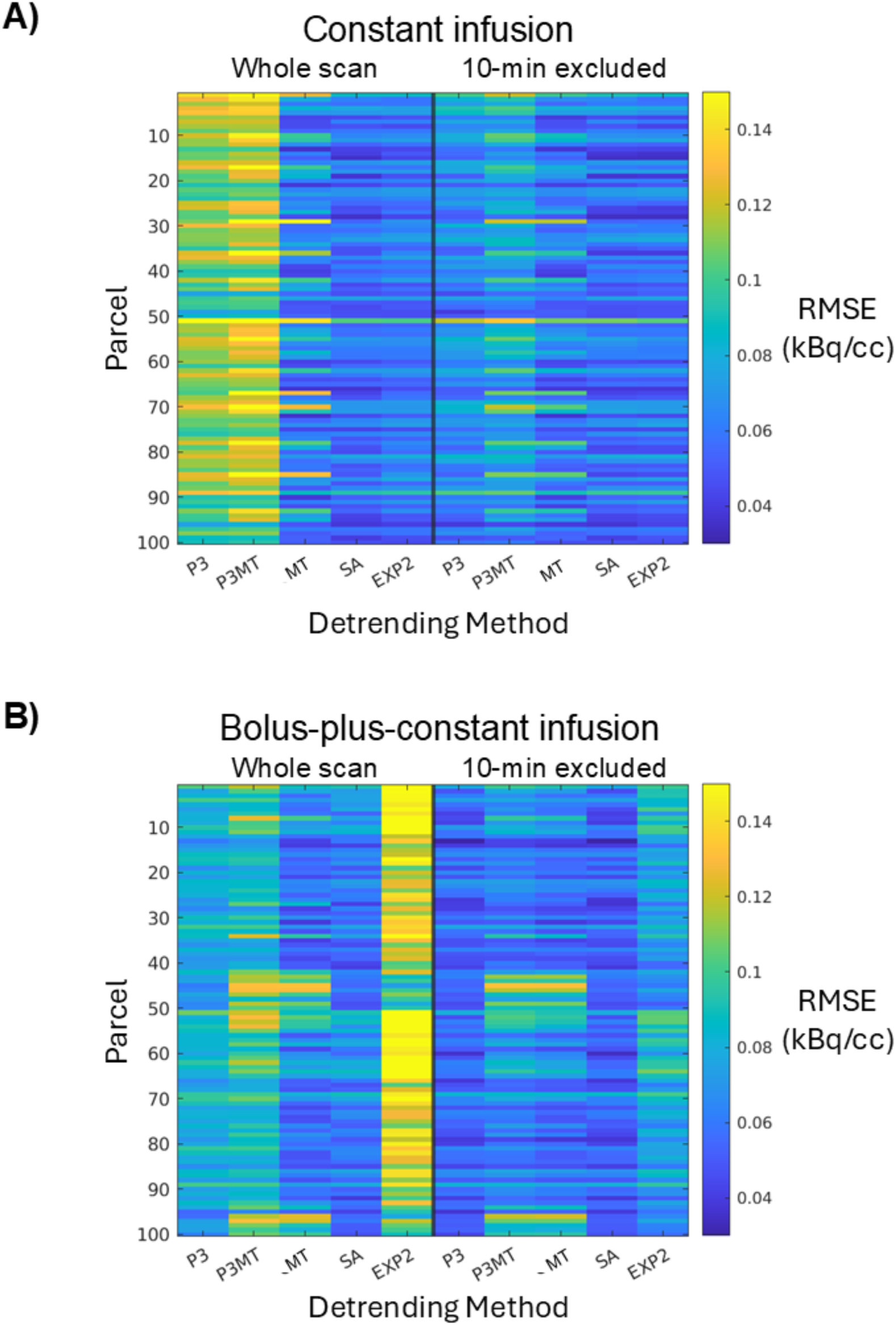
The root-mean-square error (RMSE) of the residuals in each of the 100 cortical parcels (vertical axis) from each detrending method (horizontal axis). Two-minute FWHM Gaussian temporal smoothing was applied to the residuals before the RMSE estimation to reduce the contribution from noise and thereby accentuate the contribution from residual patterning. Spatially-varying RMSEs are observed for both CI (**A**) and B+CI (**B**) paradigms.

The SA method, which requires prior knowledge of the AIF, left smaller mean residual patterns compared to P3 and P3MT for both infusion protocols (Figs. 1 and 2, “SA”). The analytical EXP2 method performed comparably to SA for the CI data but left the largest residual patterns in the B+CI data (“EXP2” vs. “SA”). This is possibly because the simplification of the AIF as comprising a single exponential failed to sufficiently model the initial dynamic phase of the true input function when appending a bolus injection.

Excluding the initial 10-minute transient phase from the analysis reduced the extent of residual patterning, particularly for polynomial detrending and in the B+CI data (“P3” and “P3MT”).

When not excluding the initial pre-equilibrium portion of each scan, a strong similarity was observed between the residuals of the simulated TACs and of the empirical data (Fig. 1, blue vs. red) for all detrending approaches except SA. This similarity suggests that the observed misfits in these cases may indeed emerge from inherent mischaracterizations of the baseline TAC shape rather than noise or unmodeled true metabolic dynamics. The simulated and empirical residuals were less similar for the approaches which excluded the first 10 minutes of the TACs, although common features can still be identified.

### 3.2 Artifactual metabolic (de)activations introduced by baseline detrending

#### 3.2.1 Artifactual effects introduced by various detrending methods

Having characterized the residual patterning resulting from various detrending approaches, we next examined whether the choice of detrending methods could introduce structured artifactual (de)activation patterns in GLM analysis in the resting-state CI dataset. Figures 3 and 4 present group-level random-effects t-scores from applying the sham regressors to the CI data, including and excluding (respectively) the initial 10 minutes. The illustrative sham regressors (Figs. 3 & 4, A) were selected based on their degrees of correlation to the mean residual pattern left by P3MT detrending (see Section 2.5). As anticipated, when detrending with a third-order polynomial, higher correlation between the task regressor and the residual pattern resulted in higher t-scores—with Regressor 1 exhibiting widespread significant (*p* < 0.05, FDR corrected) artifactual activation (Figs. 3 & 4, B, i and ii, column 1) and Regressor 3 producing some significant artifactual deactivation (Figs. 3 & 4, B, ii, column 3) for both P3 and P3MT detrending. In line with the reduced extent of residual patterning associated with excluding the initial 10 minutes observed in Section 3.1, excluding the initial period also reduced the statistical significance of the artifactual effect particularly for P3 (Fig. 3 vs. 4, B). The remaining artifactual effect after exclusion of the initial period for P3MT and MT indicates a lack of flexibility to fit various local TAC shapes as the primary driver of baseline mischaracterization for these methods, as opposed to a failure to capture early TAC dynamics.

**Figure 3:**
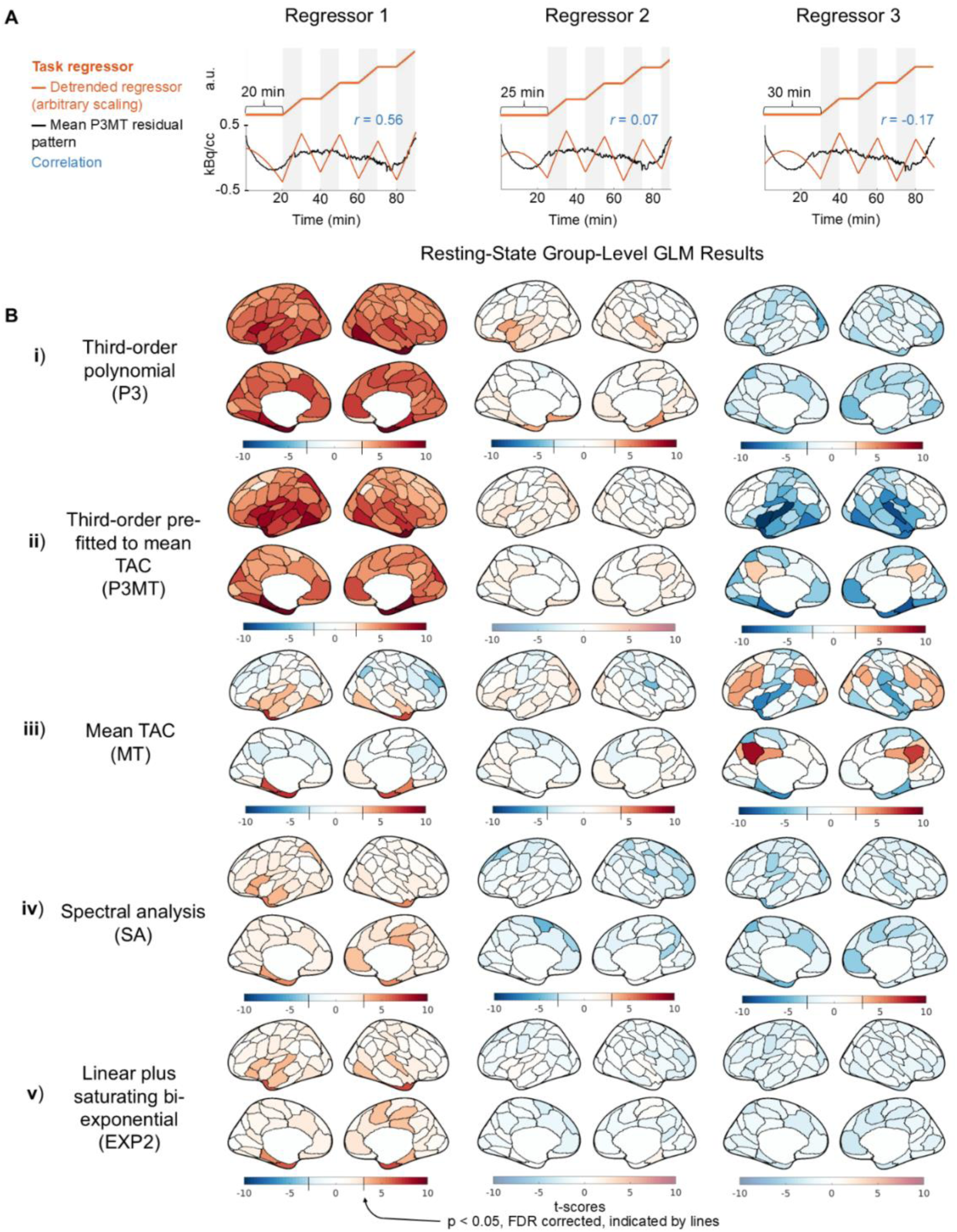
Artifactual metabolic (de)activations introduced by correlation between sham task paradigms and the residual pattern after detrending, including the entire scan time for GLM analysis for the CI data. (**A**) Sham task paradigms (orange) and their correlations (blue) with the mean residual pattern of P3MT-detrended resting-state CI TACs (black). (**B**) Group-level random-effect t-scores of the regressors applied to resting-state CI data using various baseline detrending methods (“Third-order polynomial (P3)”, “Third-order pre-fitted to the mean TAC (P3MT)”, “Mean TAC (MT)”, “Spectral analysis (SA)”, and “Linear plus bi-exponential model (EXP2)”). For the polynomial detrending methods, the significance of the artifactual effect increases with the correlation of the regressor to the residual pattern. Note that to facilitate the visualization of artifactual metabolic (de)activations, the color bar uses a step-change in saturation at the significance threshold (*p* < 0.05, FDR), if such a threshold exists (Taylor et al., 2023). Regions below the significance threshold are shown in desaturated colors, indicating insignificance, while regions above the threshold are shown in saturated colors, indicating significance. The point where the color bar transitions from desaturated to saturated varies by sub-panel, as the FDR-corrected significance threshold depends on the t-score distribution for each sub-panel.

**Figure 4:**
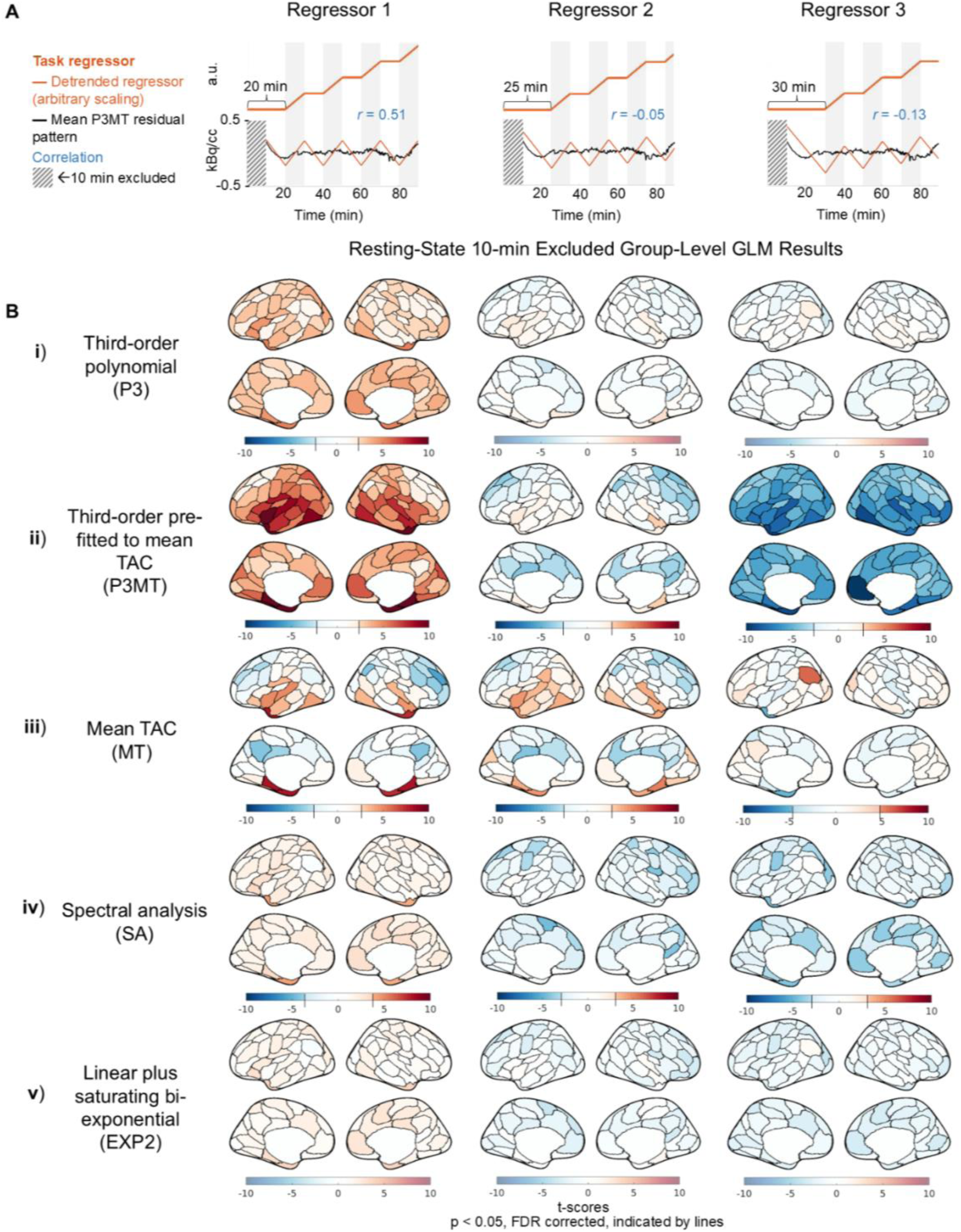
Artifactual metabolic (de)activations introduced by correlation between sham task paradigms and the residual pattern after detrending, excluding the first 10 minutes for GLM analysis of the CI data. See captions of Fig. 3 for descriptions of task regressors, detrending methods, and color scale schemes.

MT detrending resulted in regional artifactual effects (Figs. 3 and 4, iii), despite the mean effect across the brain averaging zero. The spatial pattern of these artifacts, as well as those from the other detrending methods, indicate that anatomical information is encoded in the spatial distributions of the artifactual effects, possibly due to regional differences in tracer kinetics (Heiss et al., 1984; Volpi et al., 2023).

Consistent with residual patterning results characterized in Figure 1, both SA and EXP2 produced fewer artifactual (de)activations (Figs. 3 and 4, iv and v), indicating they may improve over polynomial and mean TAC detrending. While only three shifts for the sham regressor are shown, further investigation indicates that these methods generally lead to smaller artifactual effects across all temporal shifts for the CI data (see Figure S4).

#### 3.2.2 Effect size of artifactual metabolic (de)activations

We quantified the mean absolute PSC for statistically significant (*p* < 0.05, FDR corrected) artifactual (de)activations for the 100-parcel data. The PSCs ranged from approximately 8%, the worst case observed with P3MT, down to about 2.8%, the best case observed with MT. Most regressor/detrending combinations which yielded significant artifactual effects produced mean absolute significant PSCs around 4%. While these effect sizes are smaller than those reported in previous fPET studies with robust stimuli (e.g., ∼10-30% in Godbersen et al., 2024; Hahn et al., 2016; Villien et al., 2014), the bias in estimating the task effect is consistent enough across subjects to reach statistical significance at the current sample size.

#### 3.2.3 Dependence of artifactual effects on spatial smoothing

As expected, increased levels of applied spatial smoothing were associated with higher t-scores, indicating that artifactual effects become more pronounced with greater spatial averaging; although no clear relation to spatial smoothing was observed for PSC. As an illustration, Fig. 5 displays the t-scores and PSCs for regressor 3 with P3MT detrending applied at various spatial resolutions. In Figure 5A, the average t-scores across the cerebral cortical gray matter increases from -1.2 at the voxel level to -2.3 for the 100-parcel level. Figure S5 plots t-score histograms of other sham regressor and baseline regressor combinations, showing similarly increasing trends with spatial smoothing. Still, artifactual (de)activations are observed in the voxel-wise analyses. In contrast with the trend observed in t-score magnitudes, no clear relationship was observed between PSC magnitude and spatial smoothing. In Figure 5B, the mean PSC increases in magnitude from -1.6% in the voxel-wise analysis to -2.4% at the 500-parcel level but decreases slightly to -2.2% at the 100-parcel level (see Figure S6 for PSC histograms of other sham regressor and baseline regressor combinations). The relationship between PSC and spatial smoothing may be more complex than that of t-scores for several reasons: t-scores are more influenced by noise and inter-subject variability, and smoothing may begin to decrease PSC if the smoothing area exceeds the intrinsic resolution of anatomical variations in tracer kinetics.

**Figure 5:**
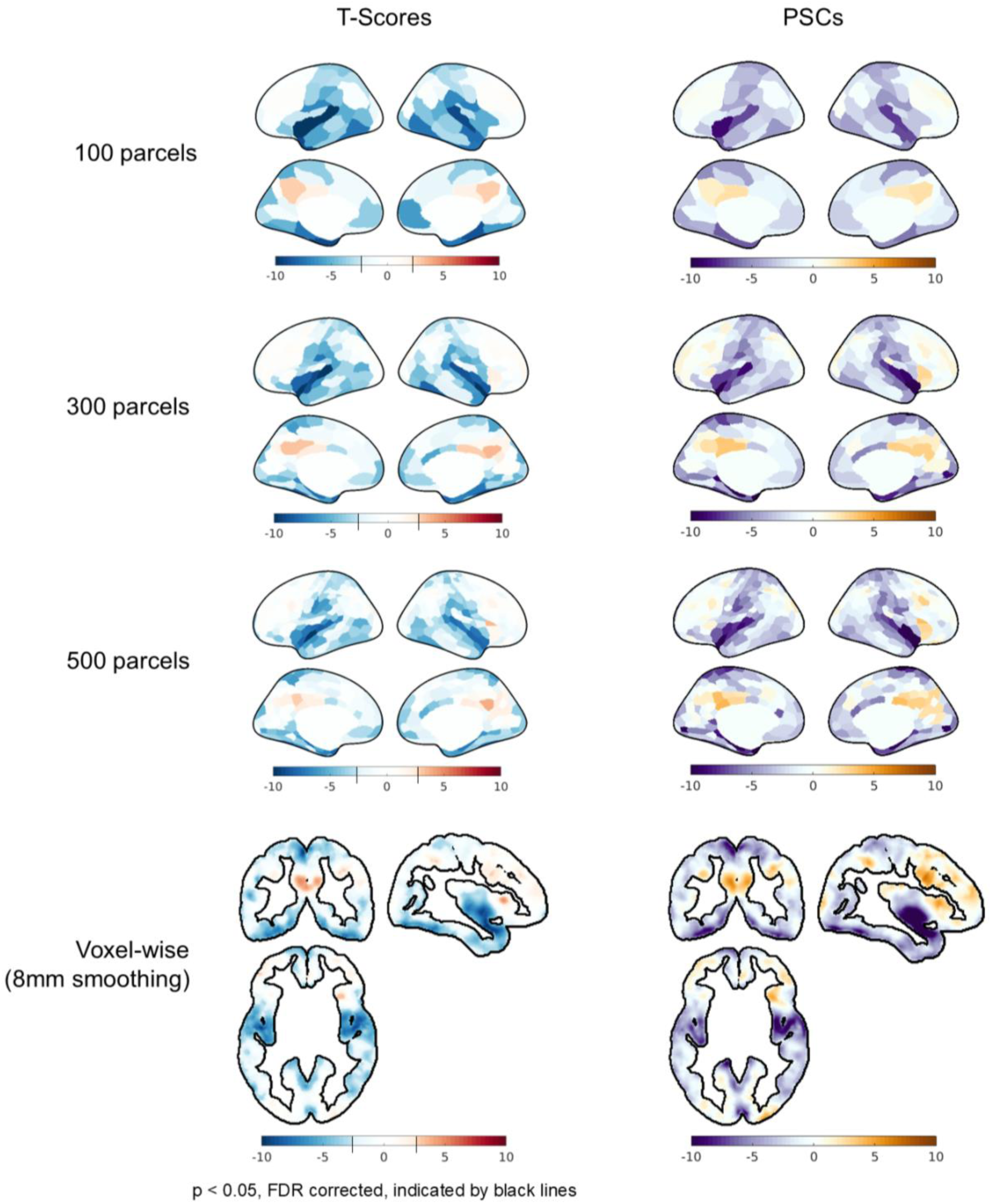
Influence of spatial smoothing (“100 parcels”, “300 parcels”, “500 parcels”, and “Voxel-wise (8mm smoothing)”) on the statistical significance (“T-scores”) and percent signal change of fPET TACs (“PSC”) of CI data associated with the artifactual effect from baseline mischaracterization. For the voxel-wise results, only cortical gray matter voxels are shown for comparison. More smoothing increases t-scores but has no discernible effects on percent signal change. Note that for the t-score plots, the color bar has a step-change in saturation at the significance threshold (*p* < 0.05, FDR), if one exists; thus the color scale varies across sub panels. The color scale representing PSC is consistent across plots.

## 4. Discussion

### 4.1 General findings

In this work, we demonstrate that mischaracterization of the baseline term of fPET-FDG TACs in GLM analysis can introduce artifactual task effects, and that the occurrence and extent of artifactual task effects depends on interactions of detrending method, infusion protocol, task paradigm, and spatial smoothing, among other factors. By combining simulations and empirical data analysis, these investigations establish that in some conditions within the realm of currently practiced methods, bias from baseline mischaracterization can result in statistically significant, anatomically coherent patterns of artifactual metabolic (de)activation.

Our findings indicate that common detrending methods like P3, P3MT, and MT can fail to accurately model the baseline [^18^F]FDG TAC shape at regional (P3MT, MT) and/or global (P3, P3MT) levels in both constant-infusion and bolus-plus-constant-infusion data (see Figures 1 and 2). A third-order polynomial was not sufficient to capture the shape of the baseline TAC in the data we analyzed, resulting in a globally consistent pattern being introduced to the residuals post-detrending. Meanwhile, based on our observation of regionally consistent patterns in the residuals of mean TAC detrending, the average TAC across the gray matter did not have the flexibility to simultaneously represent all regional TAC baselines, which vary due to anatomically informed tracer kinetics. Fitting a third-order polynomial to the mean TAC and then scaling it voxel-wise exhibits aspects of both these types of baseline mischaracterization.

Excluding the first 10 minutes of the scan substantially decreased the artifactual effects, particularly for P3 detrending, suggesting that the polynomial fails primarily to characterize the pre-equilibrium dynamics of the TAC. However, excluding the first 10 minutes did not substantially improve the performance of P3MT detrending, supporting the idea that regionally varying tracer kinetics, rather than pre-equilibrium TAC dynamics, drive mischaracterization for these methods.

The effect of these residual patterns on GLM analysis depends on their correlation with the task regressor, with the worst-cases in our dataset producing artifactual effects as large as 8 PSC, and a typical case about 4 PSC. Although this effect is smaller than those reported in task studies (10-30%), the consistency across subjects can nonetheless result in significant group-level effects with a large enough sample size (e.g., *N* = 24 in Figs. 3 and 4).

Furthermore, while we focus on GLM-based task (de)activations in this manuscript, the effect is in theory not limited to GLM analysis. For instance, there has been emerging interest in characterizing metabolic connectivity—the temporal synchrony between regional fPET-FDG dynamics over time (Jamadar et al., 2021). The baseline fPET-FDG TAC trend, if not properly removed, would also introduce bias into metabolic connectivity estimation—the patterned residuals left by detrending could create inflated correlations that obscure the global and local metabolic connectivity, the impact of which may be even stronger than task conditions due to the smaller effect size of resting-state metabolic dynamics.

### 4.2 Mitigating the impact of baseline mischaracterization

While it is appealing for reasons of ease and flexibility to minimize assumptions about the shape of the baseline TAC, our initial findings suggest that informed models can better characterize the baseline—which could possibly provide improved power and reduced bias. Spectral analysis performed well for both infusion protocols but requires accurate AIFs derived from blood sampling or image-derived approaches, which may represent a practical limitation (Veronese et al., 2016). Other informed methods, such as the analytical linear plus bi-exponential baseline (see Appendix), refer to the tissue compartment model and only require a model of the AIF. Such a model can be easily built for CI data, but not for B+CI protocols. Individual variations in the AIFs pose further challenges. Accordingly, the linear plus bi-exponential model examined in this study performed well for the CI case, but it did not perform well for the B+CI case—likely because it did not sufficiently characterize the true bolus-plus-constant-infusion AIF shape. More sophisticated analytical models (e.g., including more exponential terms) can be explored in future studies to identify a solution that generalizes more broadly to FDG administration paradigms.

As an alternative to improving baseline modeling, it is possible to modify the baseline TAC itself by excluding a portion of it or modifying the infusion protocol. Our initial results indicate that excluding the first few minutes of the scan improves the accuracy of a third-order polynomial for modeling the baseline shape, especially in the bolus-plus-constant-infusion case where the TAC quickly becomes linear. Thus, we may expect to observe mitigated impact of baseline mischaracterization in bolus-plus-constant-infusion studies which exclude the initial portion of the scan. Future research could identify the optimal bolus fraction to achieve and maintain linearity faster, possibly allowing simple polynomial baselines to effectively detrend the measured signal after the non-linear portion is removed.

In addition to more effective detrending methods or tracer administration schemes, it may also be possible to avoid bias from a study-design standpoint. Since the residual patterns exhibit certain degrees of consistency across subjects, jittering the task timing across imaging sessions would, in theory, cancel the bias at the group level. It may also be possible to de-correlate the task from the residual patterning at a subject level by using resting-state data or simulations to characterize the residual patterning, then designing a task paradigm with frequency and timing as orthogonal as feasible to that pattern.

### 4.3 Limitations

As previously noted, our study is illustrative in nature. Therefore, we only focused on the most common detrending methods and tested on a limited range of data and task protocols. Results from these analyses aimed to highlight several factors which influence the occurrence, extent, and spatial pattern of the artifactual effect that can emerge from baseline mischaracterization. How, and whether, these artifactual effects manifest in practice depends on the specific tracer administration, experimental design, and analysis methods used in each study. As such, this does not imply that artificially introduced effects are necessarily introduced in all previous studies. We also acknowledge limitations in the empirical data analyzed in this study. First, due to the extended scan duration and intermittent MR scans during the experimental sessions, the resting-state CI dataset may include endogenous or exogenous arousal and attentive effects. Second, the jittered task and random arousal effects may not entirely cancel out in the composite B+CI dataset. Therefore, the two empirical datasets used in this study may not represent perfectly task-free conditions, despite our observation that the resulting global patterns largely align with our simulations in both cases.

## 5. Conclusion

Recent studies have established fPET-FDG as a reliable technique for functional metabolic imaging. Yet, accurate baseline modeling is crucial for robust statistical analysis of fPET-FDG data. Mischaracterization of the baseline TAC can potentially introduce artifactual (de)activations to GLM analyses. Informed baseline modeling methods that incorporate prior knowledge of tracer kinetics, optimized tracer administration protocols, and careful experimental design can all help mitigate this issue. By addressing baseline mischaracterization, we can enhance the reliability of fPET-FDG in capturing true metabolic dynamics in neuroimaging research.

## Acknowledgments

This work was supported in part by the NIH (grants K99/R00-NS118120, R01-MH111438, P41-EB030006, and R21-MH135201), by the Harvard Mind Brain Behavior Faculty Research Award, by the Brain & Behavior Research Foundation Young Investigator Grant, by the BrightFocus Foundation Research Grant, by the MGH/HST Athinoula A. Martinos Center for Biomedical Imaging, and by the Northeastern Undergraduate Student Cooperative Education program; and was made possible by the resources provided by NIH Shared Instrumentation grants (S10-RR022976, S10-RR019933, S10-OD010759). Computational resources were generously provided by the Massachusetts Life Sciences Center (https://www.masslifesciences.com/).

## Appendix: fPET baseline TAC fitted by a linear plus bi-exponential model

Assume fPET-FDG kinetics follow an irreversible two-tissue compartment model, we have:

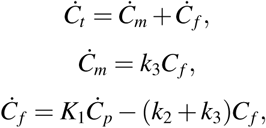

where *C_p_* indicates the plasma FDG concentration, *C_m_* indicates the metabolized FDG concentration, *C_f_* indicates the free unmetabolized FDG concentration, and *C_t_* is the sum of *C_m_* and *C_f_* .

Next, we assume a simple form of the plasma FDG concentration: *C_p_* = *a*_0_ + *a*_1_*e^-at^* (*a*_0_ and *a*_1_ are constants; note that *C_p_*is a saturating exponential function in the condition of constant-rate infusion), and let *k*_23_ = *k*_2_ + *k*_3_, then:

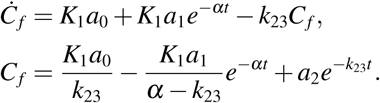

And we further have:

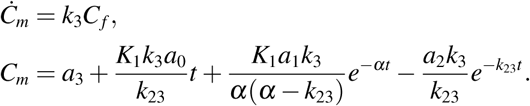

Taken together, this gives:

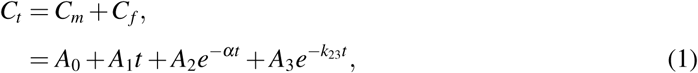

where

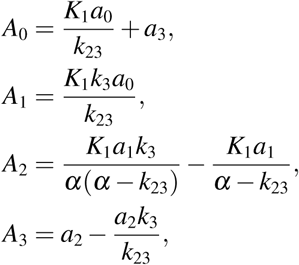

and given *C_f_* (0)= 0, *C_m_*(0)= 0, we have:

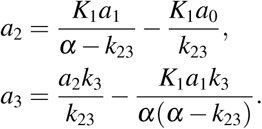

In the application of equation 1 as a baseline TAC model, we derived the exponential constants *-a* and *-k*_23_ from the mean gray-matter TAC before implementing each term in the equation as columns in the GLM design matrix. The exponential constants were estimated by first removing a linear trend from the mean TAC, fitted to the latter part of the scan (after 45 minutes), then fitting the resulting saturating biexponential directly using a method described in “Régressions et Équations Intégrales” (Jacquelin, J., *Régressions et Équations Inteǵrales*, 71-72. 2014. Scribd, https://www.scridb.com/doc/14674814/Regressions-et-equations-integrales).

