## Supplementary Figure for "On the analysis of functional PET (fPET)-FDG: baseline mischaracterization can introduce artifactual metabolic (de)activations"

### *Supplementary Figures*

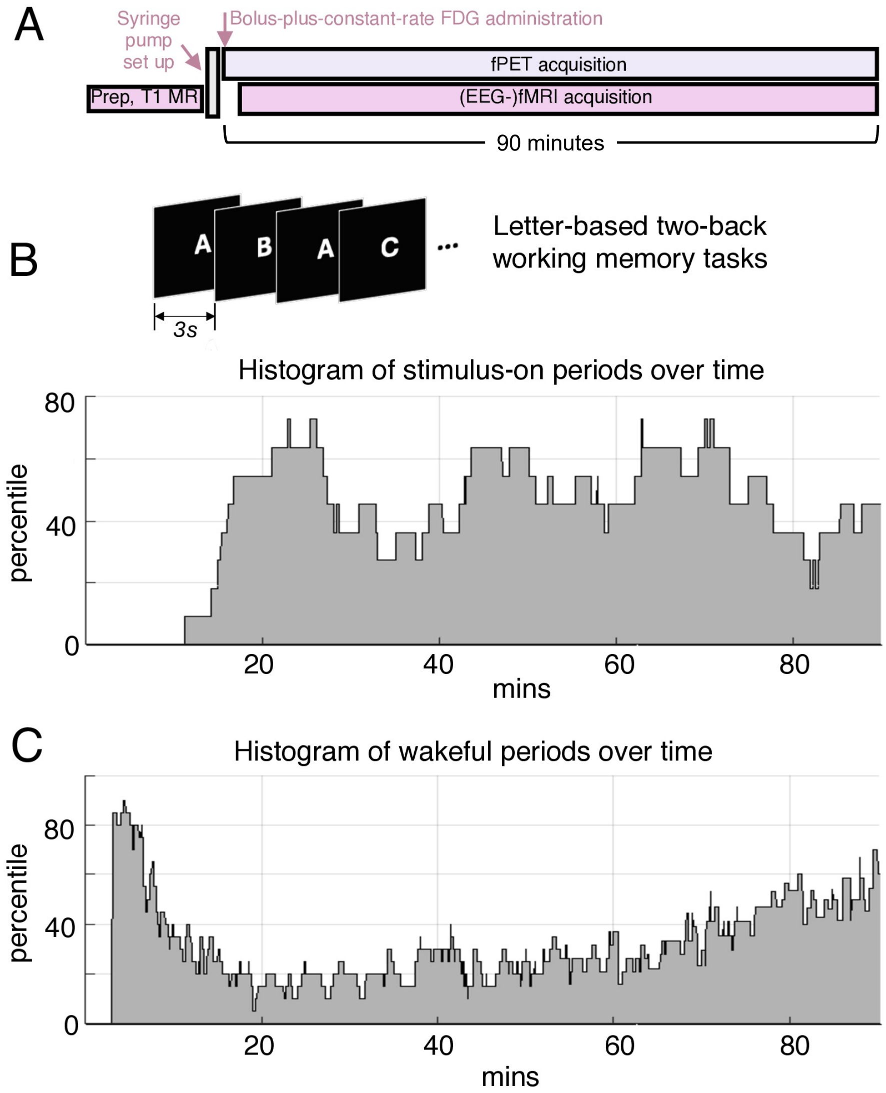


**Figure S1: Summary of the fPET-fMRI experimental schemes of the bolus-plus-constant-infusion fPET-FDG dataset, collected at MGH. (A)** Overview of the simultaneous fPET-FDG and BOLD-fMRI acquisitions. For all experiments, (EEG-)fMRI scans started a few minutes after the onset of the PET acquisition, and persisted throughout the entire experiment without interruption. **(B)** Working-memory dataset: histogram summary of the timing of stimulus-on/off blocks across 11 participants. Stimulus-on blocks (10–15 minutes): participants were instructed to judge whether a currently-present letter was identical to the one presented two letters back; stimulus-off blocks (10–15 minutes): viewing a fixation cross displayed at the center of the screen. **(C)** Endogenous-arousal dataset: histogram summary of the wakeful periods over the course of the experiment, estimated across 21 subjects with simultaneous EEG or behavioral data. This dataset was a subset of those collected to investigate sleep-induced changes in cerebral glucose metabolism in a separate study. To enhance sleep pressure during the experiment, subjects were sleep-deprived the night before the experiment, with their sleep restricted to only 4 hours.

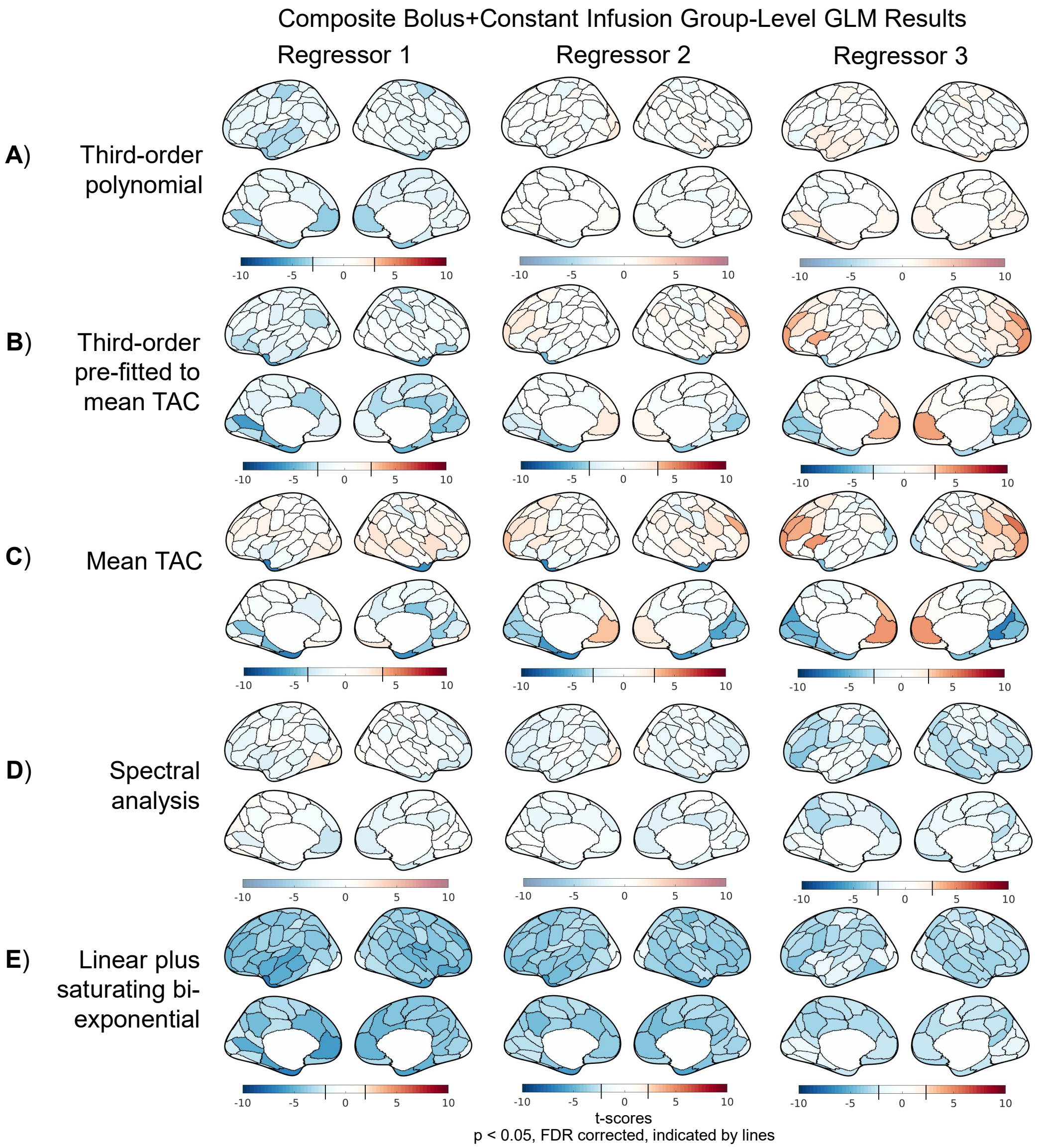


**Figure S2: Summary of group-level random-effect t-scores from applying sham task regressors to the 100-parcel composite B+CI dataset with various baseline models, including the entire scan time for GLM analysis.** The sham task regressors alternate between 10-minutes "on", modeled by a ramp with unit slope, and 10-minutes "off", modeled as flat. The sham task regressors differ in their initial rest period with Regressors 1, 2, and 3 having 20, 25, and 30 minutes of initial rest, respectively. The various baseline models comprise "Third-order polynomial (P3)", "Third-order pre-fitted to the mean TAC (P3MT)", "Mean TAC (MT)", "Spectral analysis (SA)", and "Linear plus bi-exponential model (EXP2)". To facilitate the visualization of artifactual metabolic (de)activations, the color bar uses a step-change in saturation at the significance threshold ($p < 0.05$, FDR), if such a threshold exists (Taylor et al., 2023). It should be noted that the B+CI dataset, being a composite of working memory and endogenous arousal data, may exhibit more inter-subject variability than a typical study due to between-experiment differences, complicating interpretation. Additionally, the true task and arousal effects remain present at the single-subject level, although we expect jitter and randomness to mitigate these effects to insignificance at the group level.

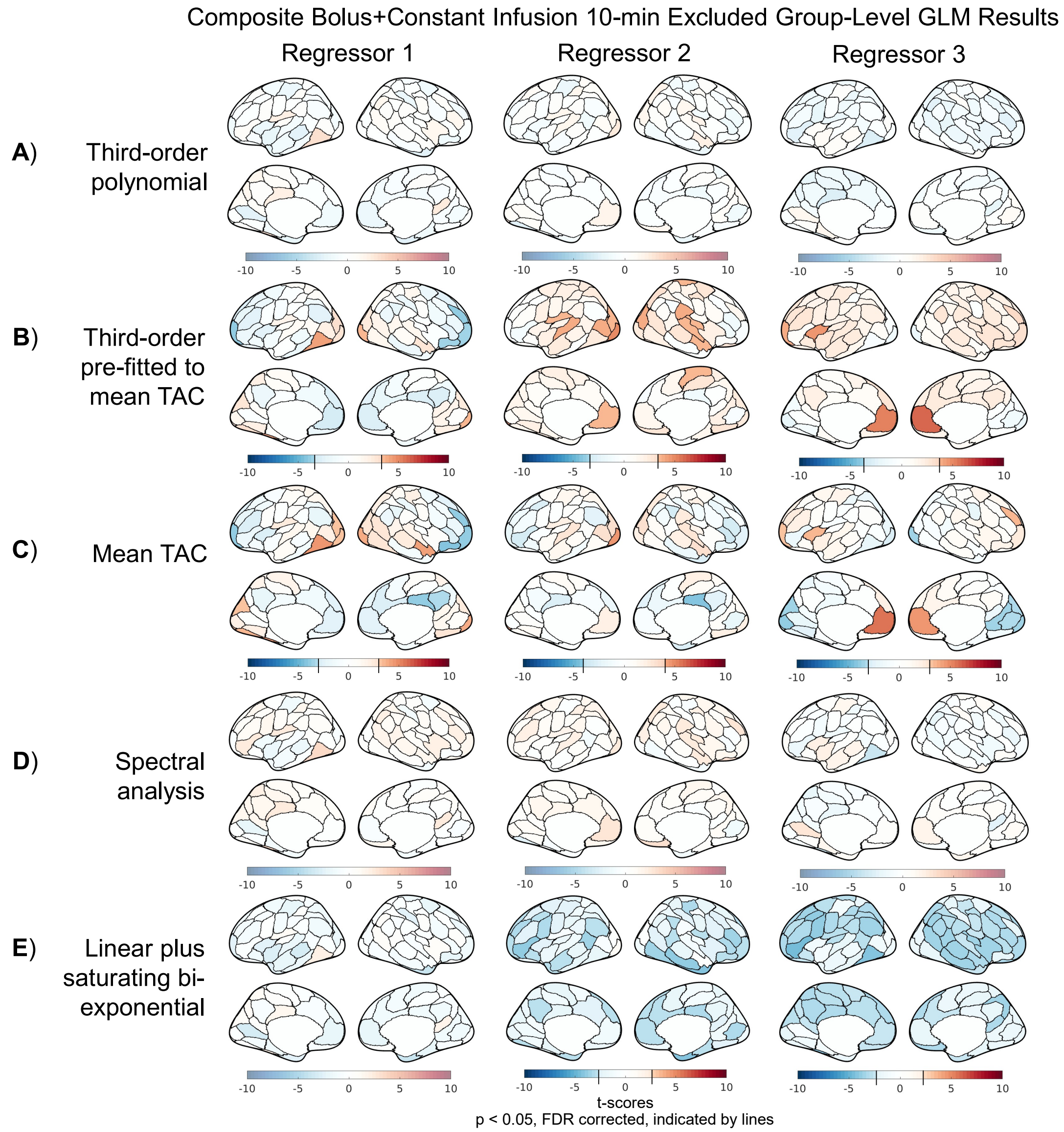


**Figure S3: Summary of group-level t-scores from applying sham task regressors to the 100-parcel composite B+CI dataset with various baseline models, excluding the first 10 minutes the scan time for GLM analysis.** Refer to the caption of Figure S2 for descriptions of task regressors, detrending methods, color scale schemes, and the B+CI dataset.

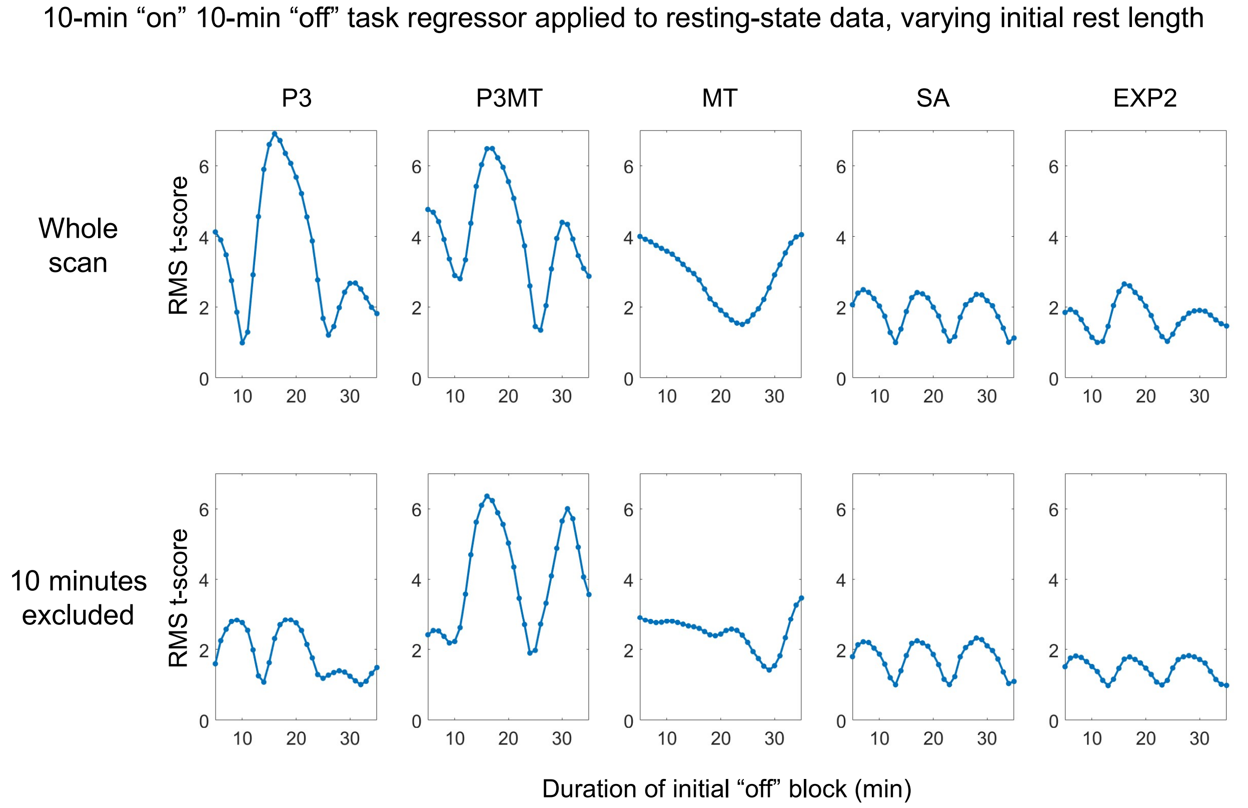


**Figure S4: Summary of group-level random-effect root-mean-squared t-scores from applying a 10-min “on”, 10-min “off” faux task regressor with varying amounts of initial rest (from 5 to 35 minutes) to the 100-parcel resting-state dataset.** Regressors 1–3, described in the main text, are the 10-, 20-, and 30-minute initial rest regressors, respectively. In the absence of an artifactual effect, the expected value of the root-mean-squared t-score across 100-parcels in 24 subjects would be 1.04, with a standard deviation of approximately 0.08.

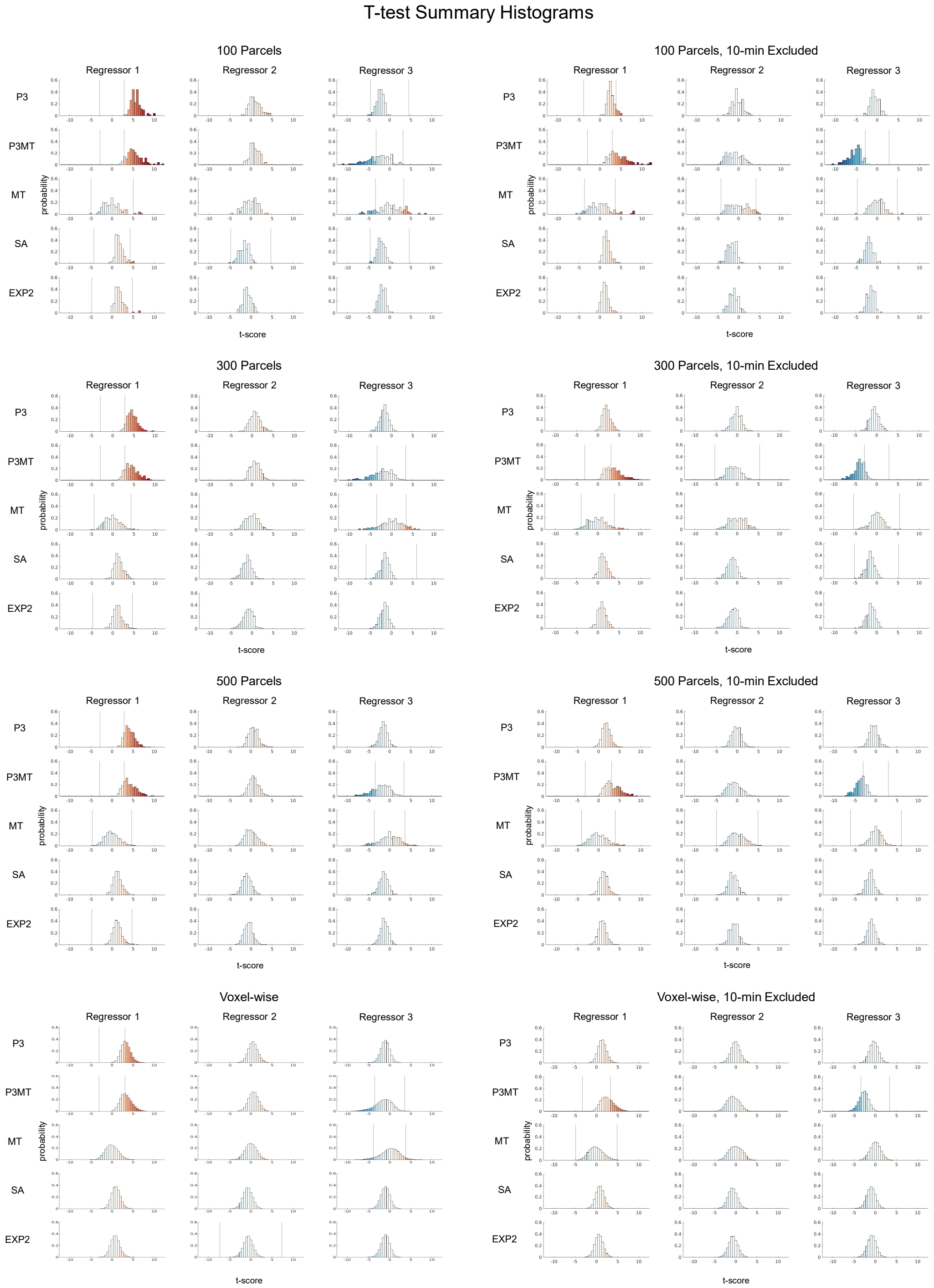


**Figure S5: Histogram summaries of t-scores for each regressor / detrending method / spatial resolution combination for the constant-infusion resting-state data.** Illustrative regressors and regression methods are identical as those used to generate results shown in Figs. 3 & 4; spatial resolutions are identical as those examined in the results shown in Fig. 5. To facilitate visualization of artifactual metabolic (de)activation, the color scale has a step-change at the ($p < 0.05$, FDR) significance point, if one exists; thus the color scale varies across sub plots.

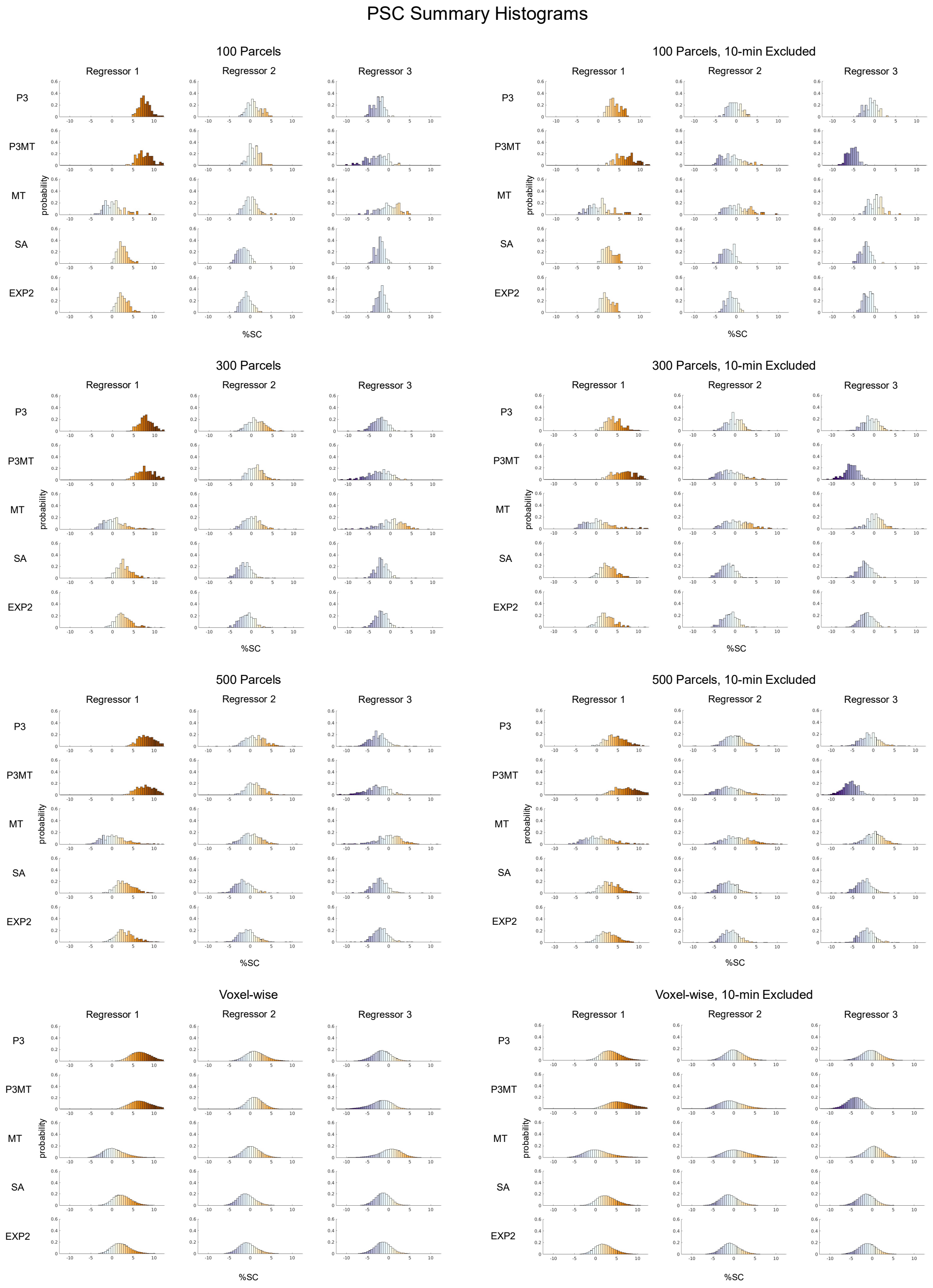


**Figure S6: Histogram summaries of percent signal changes (PSCs) of the fPET TACs for each regressor / detrending method / spatial resolution combination for the constant-infusion resting-state data.** Illustrative regressors and regression methods are identical as those used in to generate results shown in Figs. 3 & 4; spatial resolutions are identical as those examined in the results shown in Fig. 5.
